# Phenotypic sex determines recombination rate and distribution in sex-reversed Rainbow Trout *Oncorhynchus mykiss*

**DOI:** 10.1101/2025.06.04.657809

**Authors:** Cathrine Brekke, Tim Martin Knutsen

## Abstract

During meiotic cell division, homologous chromosomes align and exchange large segments of DNA through crossover recombination. Rates of recombination often show distinct differences between males and females, a phenomenon known as heterochiasmy. Despite decades of research documenting the presence of heterochiasmy across eukaryotes, the specific feature of sex leading to this curious sexual dimorphism remains to be explained. Some species, such as salmonids, also display extensive differences between the sexes in crossover positioning. A critical part of solving this puzzle is to establish whether heterochiasmy is driven by genetic sex or if it is a result of the physiological differences between producing sperm and eggs. In this study, we show that phenotypic sex determines recombination rate and distribution in hormonally sex-reversed rainbow trout. With pedigree and genotype information from 18 452 individuals and 33 913 SNP markers we map crossover events in families where the fathers were hatched as genetic XX females and sex-reversed as young trout fry with a masculinising hormone 17α-methyltestosterone and compare the crossover patterns to those in families with normal XY male fathers. We find that recombination patterns in XX males resemble those of normal XY males with crossovers exclusively in sub-telomeric regions. Crossover count per gamete was 25.8±4.4 in XX females vs 19.5±3.9 and 19.9±4.0 in XY males and XX males, respectively. These results support the hypothesis that heterochiasmy arises from physiological differences between oogenesis and spermatogenesis rather than effects related to genetic sex and will aid in guiding the research on heterochiasmy going forward.

## 1. Introduction

When germ cells are created during meiosis, homologous chromosomes align and exchange large segments of DNA through crossover recombination. This breaks up existing linkage between alleles, and together with independent assortment of chromosomes, leads to genetic variation even between gametes from the same parent. The shuffling of alleles from one generation to the next is thought to be central to the success of sexually reproducing species (Felsenstein, 1988, 1974). The process of recombination is tightly controlled, as it also has a vital functional role in the correct alignment and proper segregation of chromosomes (Fledel-Alon et al., 2009). Therefore, the number and distribution of crossovers along the chromosomes follow some shared characteristics across species such as one obligate crossover per bivalent and non-random spacing between crossovers through crossover interference (Jones and Franklin, 2006; Lenormand et al., 2016; Otto and Payseur, 2019). Despite this, recombination rates and distribution have been found to vary between individuals, populations and species (Brekke et al., 2022; C. Brekke et al., 2023; Dumont et al., 2009; Johnston, 2024; Payseur, 2024; Stapley et al., 2017). More than a century ago, Morgan discovered that there was a systematic difference in recombination rate between males and females (1914). This sexual dimorphism has since been observed across eukaryotes and ranges from the most extreme difference where only one sex recombines (Achiasmy), to the more common phenomenon where both sexes recombine but at different rates (Heterochiasmy) (Brekke et al., 2022; Broman et al., 1998; Burt et al., 1991; Isberg et al., 2006; Johnston et al., 2017; Ma et al., 2015; McAuley et al., 2023; Sakamoto et al., 2000; Sardell and Kirkpatrick, 2020). Various hypotheses have been offered to explain heterochiasmy, e.g. selection against recombination in the heterogametic sex, sperm competition, meiotic drive and sex-specific selection (Burt et al., 1991; Burt and Bell, 1987; Mank, 2009; Trivers, 1988). Yet, none of these hypotheses hold when tested empirically. A critical part of solving this puzzle is to establish whether the sex specific recombination rates and patterns are driven by genetic sex, i.e. genetic differences between males and females like sex chromosomes or sex determining genes, or if it is a result of the phenotypic sex, i.e. the sex specific morphological characteristics like ovaries and testes.

Teleost fish exhibit a broad range of sex determination systems. Their sex can be determined genetically, by environmental signals, or a combination of both (Devlin and Nagahama, 2002). In Rainbow trout, sex determination is strictly genetic, with an XX/XY system controlled by the sex-determination gene sdY (Yano et al., 2012). However, sexual differentiation in teleost fish has a liable period enabling hormonal sex-reversal (Pandian and Kirankumar, 2003). In the aquaculture industry, sex-reversal has become a common way to produce more uniform fish. After hatching, the fry is fed a diet containing 17α-methyltestosterone, causing genetic females to develop male morphological characteristics, including production of sperm upon sexual maturation. These XX males will then give exclusively female offspring when mated with normal XX females. This is favourable because all-female stock show improved uniformity of growth and reduced early onset of sexual maturation compared to mixed-sex stocks, thus improving efficiency and return for the trout farming industry (Sahafi, 2011).

Salmonids display extreme sex differences in the distribution of crossover along the chromosomes. Males recombine almost exclusively in the sub-telomeric ends of the chromosomes, whereas females recombine on a larger region of the chromosome but with a strong preference towards the centromeres (Brekke et al., 2023; Lien et al., 2011; Sakamoto et al., 2000; Timusk et al., 2011). In a previous study in Atlantic Salmon we found that both recombination rate and distribution are heritable in males and females, but with a low inter-sex genetic correlation (0.1), suggesting that the sex-specific patterns are under partly distinct genetic control and can evolve independently (Brekke et al., 2023).

The possibility for sex reversal together with the extreme sex differences in recombination makes salmonids an excellent study system to test if recombination patterns are driven by the genetic or phenotypic sex. In this study, we combine genotype data and pedigree information from 18 452 individuals from a large breeding population of Rainbow trout including more than 80 families where the fathers are XX males, i.e., individuals hatched as genetic females and hormonally treated to develop into males that produce sperm. We quantify sex-specific crossover count and crossover distribution by locating crossovers in gametes transmitted from parents to offspring. Finally, we compare crossover patterns among normal XX females, normal XY males and sex-reversed XX males and show for the first time with a large dataset of neo-males that recombination patterns are driven by the phenotypic sex and not by genetic sex.

## 2. Materials and Methods

### 2.1. Study population

The individuals in this study are from the Aquagen Rainbow trout breeding population, established in 1972 from Norwegian, Swedish, and Danish aquaculture stocks. The current breeding nucleus comprises 200 families, representing approximately 20 generations of selection primarily for large size in sea farming. Fish in this dataset were born between 2011 and 2020.

### 2.2. Sex reversal

Genetic females were sex-reversed using a standardized protocol for hormonal treatment. After hatching, young trout fry were given a feed treated with 17α-methyltestosterone at a concentration of 3 mg/kg for the first 75 days of feeding. Putative neo-males were tested using DNA-based sex determination assays to confirm that their genetic sex was XX. Only individuals with successful sex reversal were included in the study. Neo-males were subsequently used as sires in controlled breeding with normal XX females to produce all-female offspring. Fertilization was performed using milt from confirmed Neo-males, following standard artificial breeding protocols.

### 2.3. Genotype and pedigree data

All fish in the study, a total of 18 452 individuals, were genotyped using the Aquagen custom Affymetrix genotyping array containing 50 000 Single Nucleotide Polymorphisms (SNPs) segregating in the breeding population and evenly distributed along the genome. Genotype calls were generated using the Thermo Fisher Best Practices Genotyping Analysis Workflow. Markers in the categories PolyHighResolution and NoMinorHom were kept for further analysis, i.e, SNPs with well-separated genotype clusters and two or more alleles in the genotype calls, and where one cluster is homozygous, and one is heterozygous for biallelic SNPs and only one homozygous cluster and one or more heterozygous clusters appear for multiallelic SNPs (Fisher Scientific, 2023). In further quality control with PLINK (Chang et al., 2015) we removed SNPs with a minor allele frequency < 0.01, genotype call rate < 0.95, and strong deviation from Hardy Weinberg equilibrium 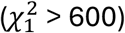. A final set of 33,912 SNP markers were used in further analysis.

### 2.4. Three generation families for phasing and crossover detection

Crossover events were mapped by detecting new haplotype combinations in an offspring compared to its parents. This was done by sorting the pedigree into multiple three-generation full-sib families in the following manner: Each unique mating pair made the basis of a family with its offspring and, if known, their parents. This structure allows for genotype phasing of the offspring and mating pair and detection of crossovers in the offspring gametes that occurred during meiosis in the parents. An individual can be in more than one family (e.g. as a parent in one and an offspring in another one), but each meiotic event is only counted once. For parents where grandparents were missing, we kept only families that had at least four offspring and where the grandparents of the mates was known, to ensure proper phasing. This resulted in 53 unique families in the pedigree with normal XY male fathers and normal XX female mothers and 80 unique families in the pedigree with sex-reversed XX male fathers and normal XX female mothers. The number of offspring in each family ranged from 4 to 198.

### 2.5. Sex specific linkage mapping and individual crossover detection

Phasing and linkage mapping were performed using Lepmap3 (Rastas, 2017). SNP marker physical positions were first mapped to the USDA_OmykA_1.1 Rainbow trout genome. In Lepmap3, the *filtering2* module was run as suggested for multi-family datasets with a *datatolerance* = 0.01 to filter markers based on segregation distortion. The *separatechromosomes2* module was run within linkage groups and markers that were not assigned to the main group (LOD score < 5) were excluded, as suggested in species where chromosome-level assemblies and marker positions are well established. The *ordermarkers2* module was run with the option to evaluate the given marker order, i.e., to calculate the centimorgan (cM) positions using the Morgan mapping function option. Sex specific recombination rates were estimated for two different datasets, one with the families with normal XY fathers and one with the families with XX fathers, allowing to compare recombination patterns for XX males, XY males and XX females separately. Final XX female recombination rates were estimated based on XX females from both datasets. Crossover counts were determined for each gamete by counting crossovers from the phased output of the *orderMarkers2* module, the total number of crossovers across chromosomes from a single gamete was assigned as an observation to the parent where meiosis took place. Gametes with no crossovers on any chromosomes or with a total crossover count higher than 50 across chromosomes were excluded from further analysis. This resulted in a dataset with crossover observations from a total of 8 945 gametes, 1 691 gametes from 30 XY males, 2 782 gametes from 44 XX neo-males and 4 472 gametes from 91 XX females. Differences in crossover count between the three groups were tested with the Welch t-test (Welch, 1947).

## 3. Results

### 3.1. Sex-specific individual crossover counts

Mean genome-wide crossover count in gametes from sex-reversed XX males was 19.9 (±4.0), similar to that of normal XY males, with crossover count = 19.5 (±3.9). Crossover count in gametes from XX females was significantly (p = 2.2×10−16) higher, with mean 25.8 (±4.4) (Figure 2a and Table 1). In all three groups this is consistent with at least one obligate crossover per bivalent during meiosis. Crossover count was close to normally distributed in all three groups and ranged between 10-49 in females, 9-39 in XX neo-males and 8-45 in XY males. The female maps are longer than both male maps on all chromosomes, except chromosome 5, 14, 15 and 22, where male maps are slightly longer or similar. On chromosomes 13, 23 and 26, we detected very few crossover events in both XY males and XX neo-males, possibly due to lower marker coverage in the telomeric regions on these chromosomes where male crossovers occur. Chromosome-level results from the linkage mapping for all chromosomes are provided in Table S1.

**Table 1.**
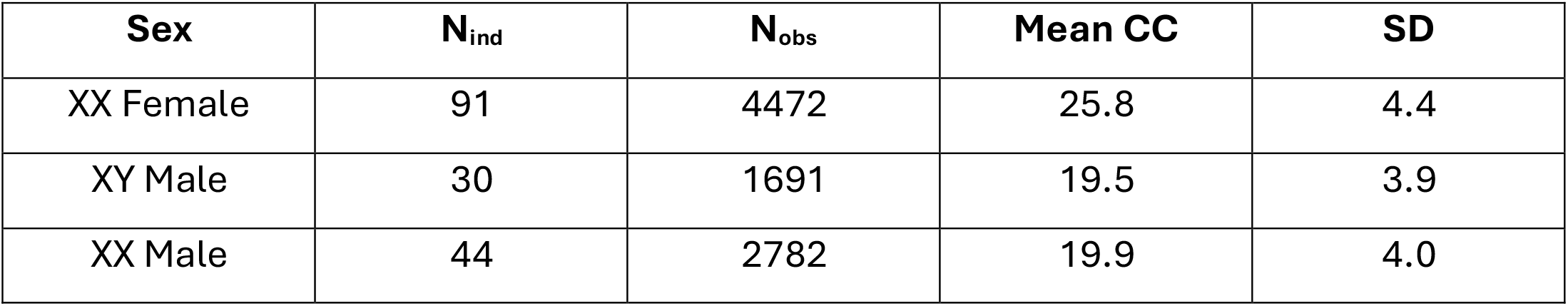
Sex-specific individual crossover counts. N_ind_ is the number of unique parent individuals and N_obs_ is the number of observations, i.e. the total number of gametes from each sex. Mean CC is the mean crossover count for each sex and SD is the standard deviation.

**Figure 1.**
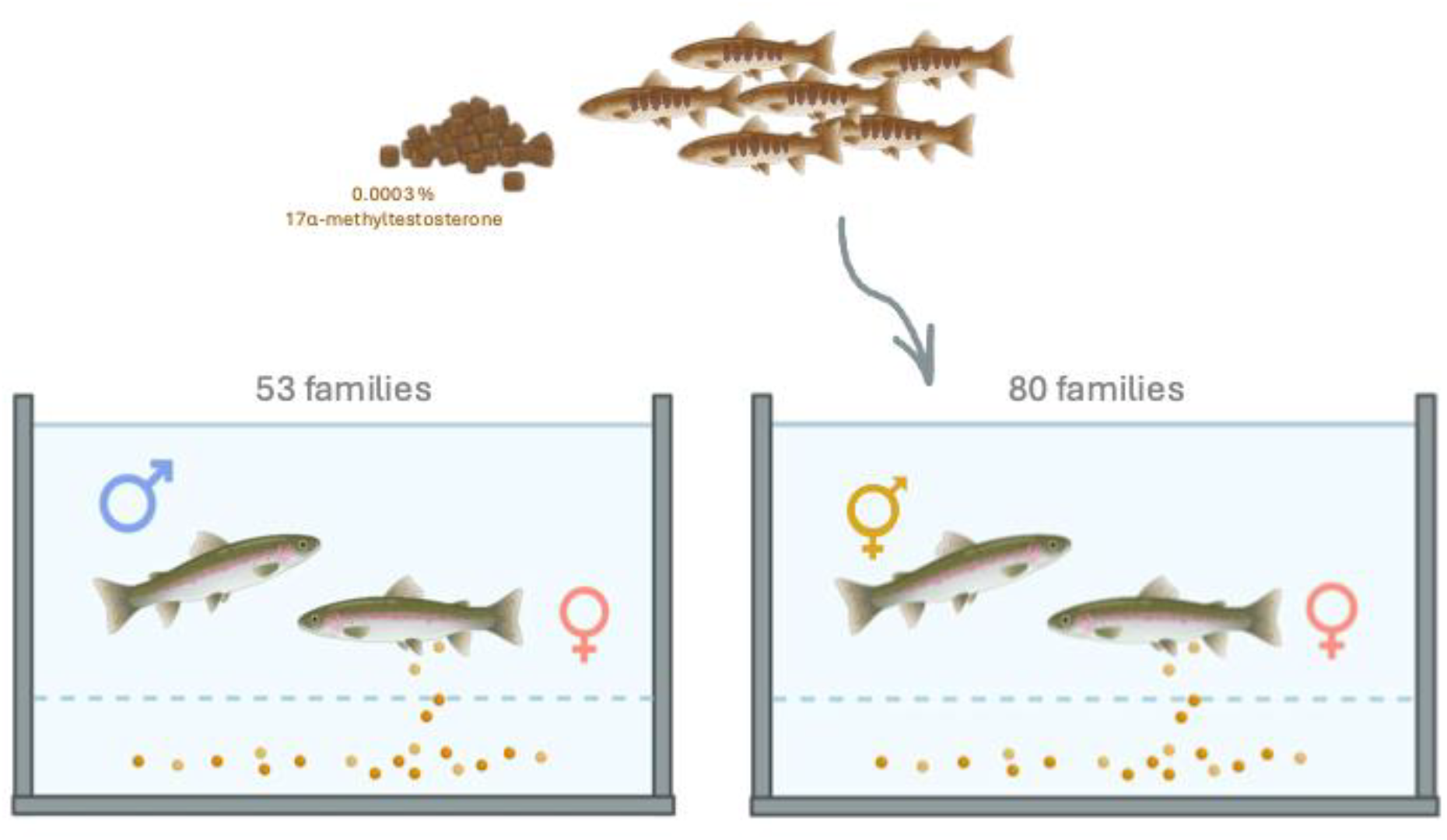
Study design. Young Rainbow trout fry were sex reversed by adding 17α-methyltestosterone to the feed. After sexual maturation normal XY males (left) and sex-reversed XX males (right) were mated with normal XX females. Parents and several offspring from each family were genotyped for phasing, linkage mapping and crossover detection.

**Figure 2.**
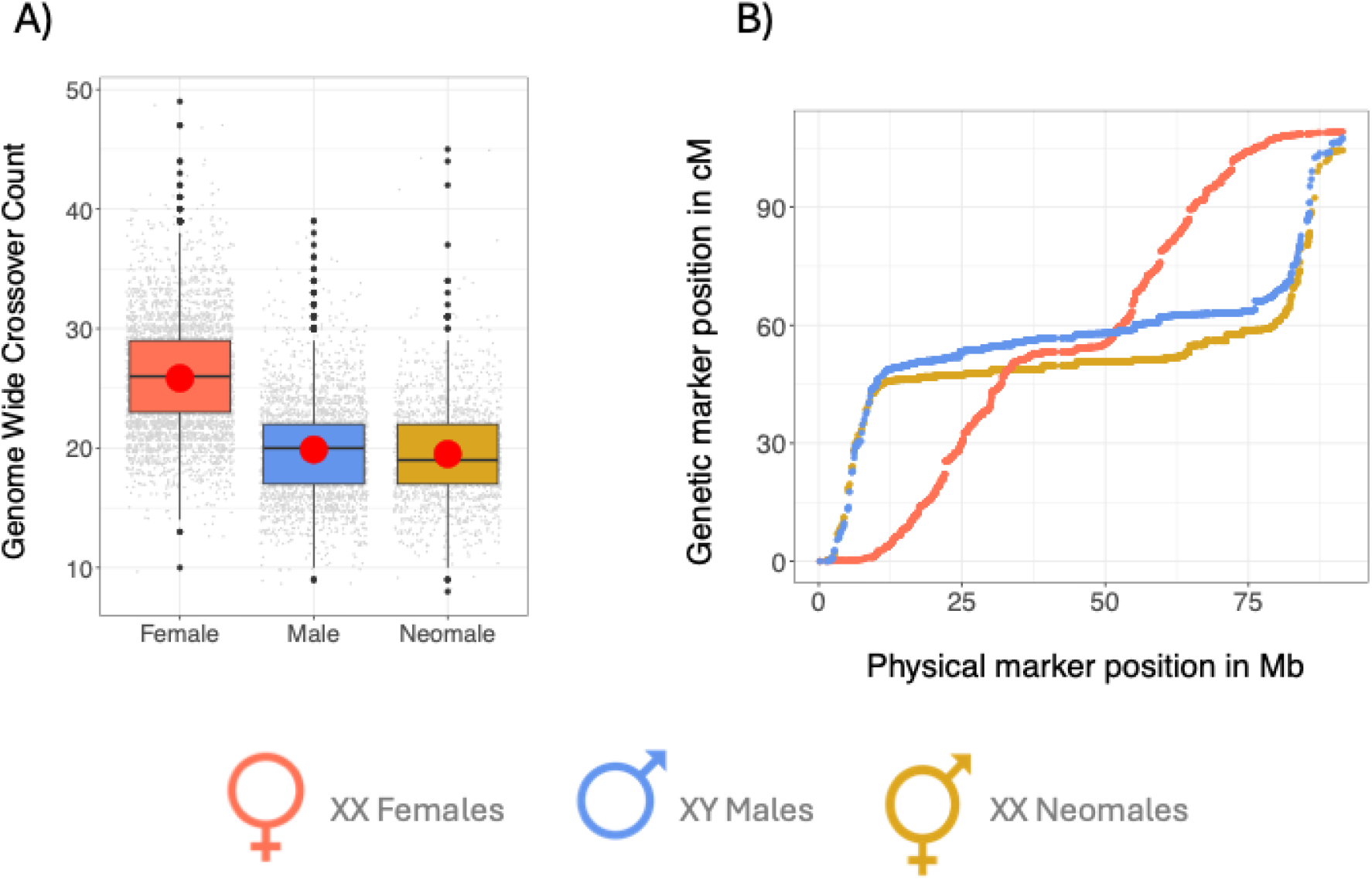
Sex-specific crossover count and distribution. A) comparison of crossover counts in XX females (red), XY males (blue) and XX neo-males (yellow). The red dots are the means, the midline in the boxplot is the median and the box is from the 25th percentile to the 75th percentile. All observations for the three respective groups are plotted in grey scatter. B) Marey map of the metacentric Rainbow trout chromosome 8. Genetic positions in cM on the y-axis is plotted against physical positions in megabases (Mb) for all SNP markers on chromosome 8 on the x-axis. Marey maps for all chromosomes can be found in Supplementary Figure S1.

### 3.2. Crossover distribution

Crossover patterns in XX neo-males resembled that of XY males with crossovers almost exclusively in sub-telomeric regions, as opposed to in females where crossovers were more evenly distributed along the chromosome, and rare in sub-telomeric regions (Figure 2b and Figure S1). Recombination rates were higher on metacentric chromosomes than acrocentric chromosomes for all three groups (Figure S1), consistent with the observation that there is often at least one crossover per chromosome arm. Marey maps showing the relationship between the physical and genetic length of all chromosomes are shown in Figure S1. The full linkage map can be found in Supplementary Table S2.

## Discussion

In this study, we demonstrate for the first time in sex-reversed males that crossover rate and distribution is controlled by the phenotypic sex and not the genetic sex. By combining pedigree and genotype data from more than 130 families and 18 000 individuals in a rainbow trout breeding population we were able to accurately map crossover events occurring during meiosis in individuals that are genetically XX females but display male morphologies, including sperm production. We then compared the crossover patterns to crossover patterns in genetic and phenotypic female individuals and genetic and phenotypic male individuals and found that both the number of crossovers and the crossover patterning in XX males resemble that of XY males with crossovers occurring exclusively in the telomeric regions. This clearly contrasted the XX female recombination patterns that were more evenly distributed along the chromosomes and with a significantly higher overall crossover count than males. Here, we explore the results in more detail and discuss how the findings may be relevant for Aquaculture, as well as how they contribute to the understanding and future research on heterochiasmy.

### 4.1. Heterochiasmy is likely rooted in mechanistic differences between oogenesis and spermatogenesis

Our findings support the emerging hypothesis that heterochiasmy is not related to genetic sex but rather caused by mechanistic differences between oogenesis and spermatogenesis. Our results support the studies by Wallace et al., 1997, Kondo et al., 2001 and Lynn et al., 2005 that found that sex-reversed females displayed the same recombination patterns as normal females in mice, crested newts and medaka, respectively. These studies were limited to females and examined only a small number of meiotic events and a subset of chromosomes. Our study confirms these findings in sex-reversed males, includes all the rainbow trout chromosomes and is based on observations from more than 8 000 meiotic events.

Oogenesis and spermatogenesis display several known mechanistic differences in the meiotic cell cycle. The most obvious one is that in most species the meiotic cell cycle occurs at different life stages in males and females, e.g., in mammals where female meiosis occurs already during embryo development in utero, whereas male meiosis starts at puberty and lasts throughout life. Potential differences related to life stages could be gene regulation and thereby chromatin accessibility for the recombination machinery at the time of meiosis (Guo et al., 2017). Even when meiosis occurs at more similar life stages in males and females, it is well established that each process of oogenesis takes longer to complete, and that the checkpoints that detect DNA damage and mis-segregation of chromosomes are not as stringent as in spermatogenesis (Cahoon and Libuda, 2019; Morelli and Cohen, 2005). It has been hypothesized that this may allow female meiosis to better cope with challenges such as a duplicated genome, due to its greater flexibility and the extended time available to resolve entanglements, multivalents, and other complex chromosomal configurations (Bomblies, 2023). This would be an interesting avenue to explore in understanding heterochiasmy in salmonids as the genomes still show signs of delayed rediploidization after whole genome duplication (Berthelot et al., 2014; Lien et al., 2016). The physical size of the chromosomes during meiosis, i.e. the length of the synaptonemal complex in μ*m* is also different in males and females across species and this difference has been found to correlate well with the direction of heterochiasmy so that the sex displaying longer chromosome axes also has higher recombination rates (Cahoon and Libuda, 2019; Gruhn et al., 2013). Recombination is known to play a role in the correct coupling of homologous chromosomes early in prophase 1, but homolog pairing and synapsis initiation can also vary greatly between species (Zickler and Kleckner, 2015). The underlying mechanisms governing these events are not yet well understood, and consequently, little is known about their sex-specific properties. For an extensive review of sex-specific differences in meiosis prophase I across model species, see Cahoon and Libuda (2019).

### 4.2. Implications for aquaculture

The crossover rates and patterns of the sex-reversed rainbow trout individuals will affect to what extent the alleles of the all-female stock are broken up and recoupled by recombination. The finding that recombination patterns in sex-reversed individuals follow their phenotypic sex suggests that all-female stock will display the same level of shuffling caused by recombination as a normal mixed sex stock from XX and XY parents.

A potential application of our findings in aquaculture could be to use sex reversal to alter recombination patterns within the breeding nucleus. Since salmonid males only recombine in a limited region at the end of the chromosomes, and the sdY allele is only passed down from father to son, unfavourable alleles on this chromosome in linkage with the sdY allele cannot be selected against. If breeding males were sex-reversed to XY females that produce eggs upon sexual maturation, it would be possible to break up linkage built up between unfavourable alleles and the sdY allele. Such an approach could significantly enhance the efficiency of selective breeding programs by increasing the shuffling of genetic material in specific parental lines.

### 4.3. Conclusion and future directions

In this study we confirm that sex-specific recombination patterns are not linked to genetic sex, but to the mechanistic differences between oogenesis and spermatogenesis.

Consistent with previous studies, we detect extreme differences in crossover distribution in a salmonid species, making them a promising study system to better understand this widespread phenomenon. Future studies should aim to compare sex-specific properties of meiotic prophase 1, particularly in mechanisms involved in axes formation and synapsis initiation and whether these sex differences are conserved or vary across species. In salmonids specifically, an interesting unexplored question is how the ongoing process of genome rediploidization after ancient whole-genome duplication relates to the extreme sex-specific recombination patterns.

## Acknowledgements

We want to thank Aquagen for providing the data for this study. Special thanks to Jørgen Ødegård, Yukiko Imai and Corinne Grey, for helpful discussions. We thank Helle Tessand Baalsrud for reading and commenting on the final version of the manuscript. Most of the analyses in this study have been executed on the Orion cluster at NMBU (https://orion.nmbu.no).

## CRediT authorship contribution statement

### Cathrine Brekke

Conceptualization, Formal analysis, Investigation, Methodology, Visualization, Writing-original draft. **Tim Martin Knutsen:** Conceptualization, Data curation, Investigation, Methodology, Writing-review and editing.

### Competing interests

Tim Martin Knutsen is employed by Aquagen. The authors declare no competing interests.

## Supplementary

**Table S1.** Sex spesific physical and genetic lengths of all chromosomes.

| Chr | physical length(bp) | Female cM | Male cM | Neo-male cM |
| --- | --- | --- | --- | --- |
| 1 | 95.77 | 111.31 | 77.11 | 64.94 |
| 2 | 103.81 | 117.96 | 88.01 | 76.64 |
| 3 | 85.31 | 109.90 | 77.06 | 60.00 |
| 4 | 46.84 | 55.53 | 51.47 | 48.52 |
| 5 | 100.80 | 125.62 | 122.65 | 128.36 |
| 6 | 101.10 | 128.99 | 102.04 | 86.83 |
| 7 | 90.92 | 117.95 | 89.21 | 83.08 |
| 8 | 91.62 | 109.19 | 107.63 | 104.43 |
| 9 | 79.46 | 109.67 | 107.01 | 90.40 |
| 10 | 87.81 | 110.32 | 98.39 | 91.65 |
| 11 | 86.28 | 109.42 | 111.36 | 108.52 |
| 12 | 102.85 | 119.43 | 84.40 | 73.77 |
| 13 | 73.33 | 89.48 | 37.99 | 19.30 |
| 14 | 43.31 | 55.32 | 56.20 | 54.77 |
| 15 | 81.57 | 102.49 | 74.80 | 61.59 |
| 16 | 78.54 | 106.79 | 113.86 | 103.25 |
| 17 | 95.21 | 104.08 | 73.18 | 59.04 |
| 18 | 74.66 | 97.97 | 64.65 | 52.44 |
| 19 | 67.24 | 84.71 | 56.01 | 55.07 |
| 20 | 46.62 | 58.44 | 57.35 | 57.37 |
| 21 | 64.94 | 83.48 | 55.50 | 47.37 |
| 22 | 52.47 | 79.07 | 101.25 | 97.52 |
| 23 | 62.88 | 59.89 | 17.86 | 8.65 |
| 24 | 45.93 | 58.66 | 67.86 | 83.77 |
| 25 | 47.54 | 59.54 | 70.26 | 48.45 |
| 26 | 51.11 | 48.79 | 4.11 | 3.69 |
| 27 | 51.56 | 53.30 | 55.22 | 47.51 |
| 28 | 43.72 | 60.44 | 57.14 | 58.86 |
| 30 | 46.33 | 54.83 | 55.54 | 50.61 |
| 31 | 44.11 | 52.05 | 51.69 | 49.92 |
| 32 | 41.84 | 57.63 | 56.14 | 53.28 |
| Y | 47.75 | 53.17 | 55.92 | 50.81 |
| sum | 2,233.22 | 2,745.40 | 2,298.84 | 2,080.39 |

**Figure S1.**
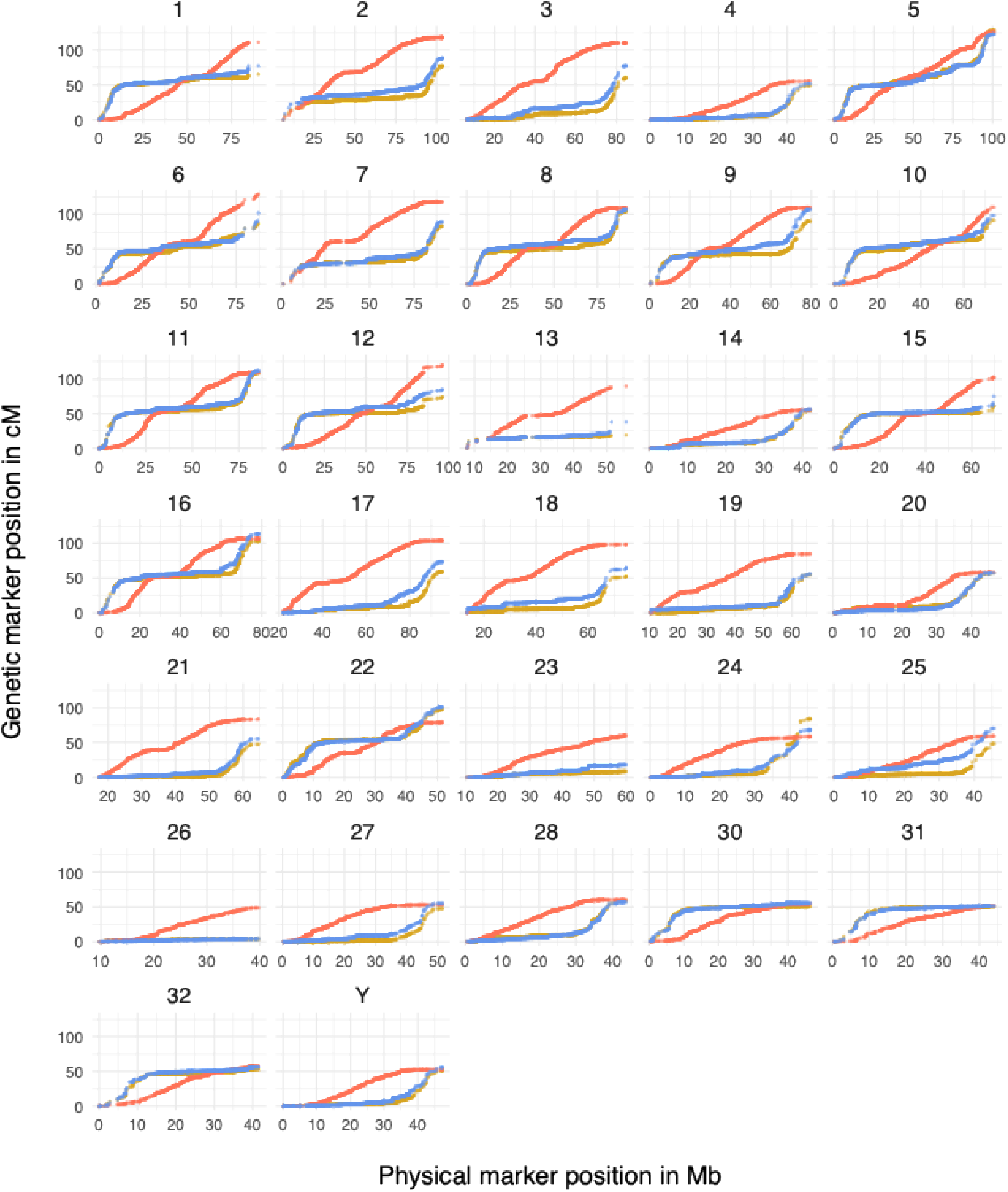
Marey map. Sex spesific Marey maps for the Rainbow trout chromosomes. The genetic position in cM on the y-axis is plotted against the physical position in Mb on the x-axis for the SNP markers within each linkage group. Female positions in red and male in blue. Maps for XX females is plotted in red, XY males in blue and XX neo-males in yellow.

## Notes

### Competing Interest Statement

The authors have declared no competing interest.

